# Generation, selection and transcriptomic profiling of human neuromesodermal and spinal cord progenitors in vitro

**DOI:** 10.1101/182279

**Authors:** Laure Verrier, Lindsay Davidson, Marek Gierliński, Kate G. Storey

## Abstract

Robust protocols for directed differentiation of human pluripotent cells are needed to establish the extent to which mechanisms operating in model organisms are relevant to our own development. Recent work in vertebrate embryos has identified neuromesodermal progenitors as a bipotent cell population that contributes to paraxial mesoderm and spinal cord. However, precise protocols for *in vitro* differentiation of human neuromesodermal progenitors are lacking. Informed by signalling activities during spinal cord generation in amniote embryos, we show here that transient dual-SMAD inhibition, together with retinoic acid (dSMADi-RA), provides rapid and reproducible induction of human spinal cord progenitors from neuromesodermal progenitors. We use CRISPR-Cas9 to engineer a GFP-reporter for a neuromesodermal progenitor-associated transcription factor *Nkx1.2* in human embryonic stem cells, to facilitate selection of this cell population. RNA-sequencing (RNA-Seq) was then used to identify human and conserved neuromesodermal progenitor transcriptional signatures, validate this differentiation protocol and implicate new pathways and processes in human neural differentiation. This optimised protocol, novel reporter line and transcriptomic data are useful resources with which to dissect cellular and molecular mechanisms regulating the generation of human spinal cord, allow scale-up of distinct cell populations for global analyses, including proteomic, biochemical and chromatin interrogation and open up translational opportunities.

## Introduction

Head and trunk nervous systems have distinct developmental origins. Head or anterior neural progenitors are derived from the epiblast rostral to the primitive streak and will form regions of the brain. In contrast, progenitors of trunk or posterior neural tissue (posterior hindbrain and spinal cord) arise from epiblast adjacent to and within the anterior primitive streak (known as caudal lateral epiblast (CLE) and node streak border (NSB), respectively) (Wilson et al. 2009) (Fig. 1A). In recent years, evidence has accrued which indicates that, unlike anterior, posterior neural tissue is generated via an intermediary neuromesodermal progenitor (NMP), which contributes paraxial mesoderm as well as posterior neural tube (reviewed (Tzouanacou et al. 2009; Gouti et al. 2015; Henrique et al. 2015; Tsakiridis and Wilson 2015). Human, mouse and chick embryos as well as *in vitro* NMPs are identified by co-expression of early neural (Sox2) and mesodermal Brachyury (Bra) proteins, but as yet lack unique molecular markers (Olivera-Martinez et al. 2012; Gouti et al. 2014; Turner et al. 2014; Henrique et al. 2015; Tsakiridis and Wilson 2015). While we are beginning to uncover how mouse NMPs are regulated, human NMPs and their derivatives are less well characterised, in part because this requires creation of robust in vitro models.

**Figure 1.**
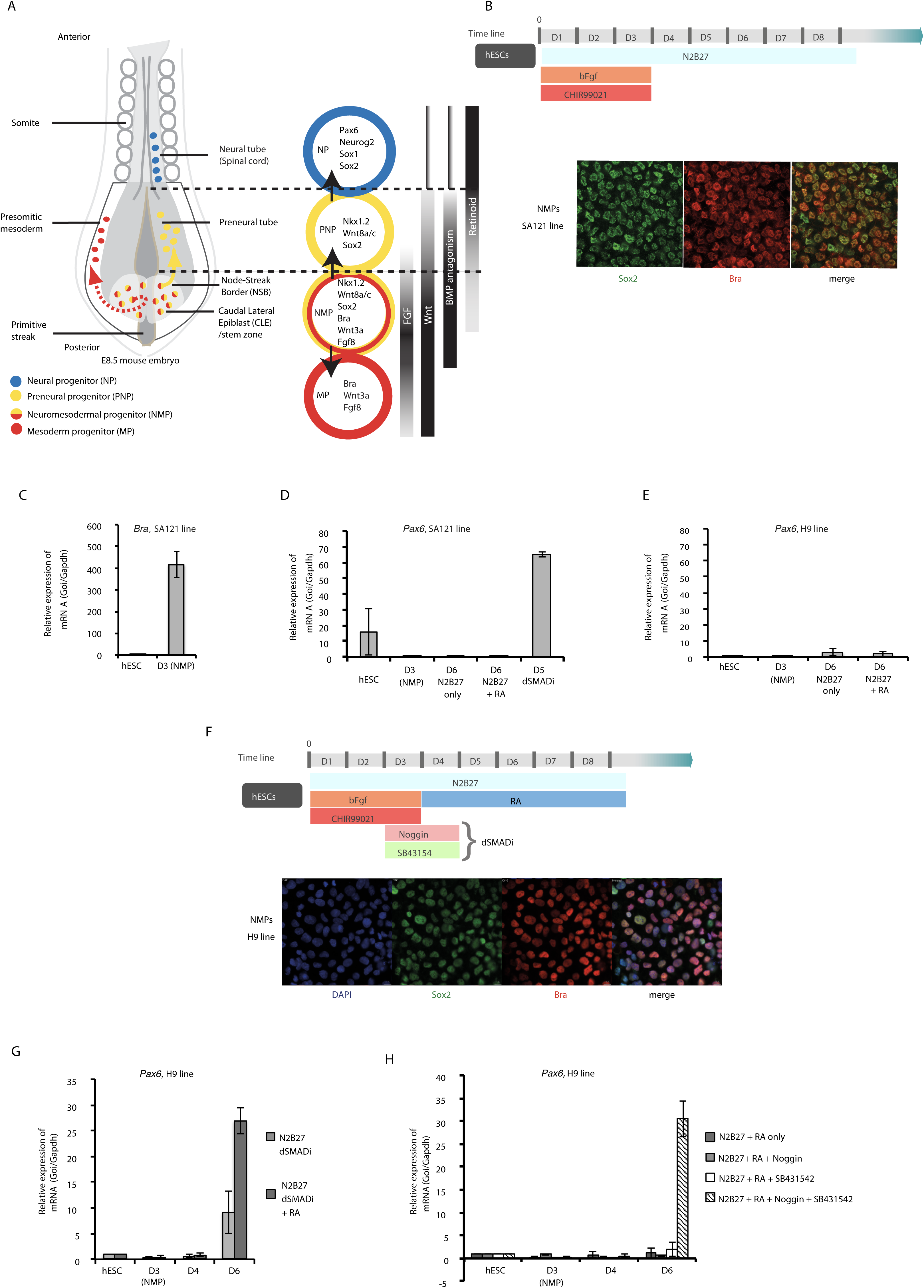
Protocol for neural differentiation of human neuromesodermal progenitors. A. Schematic of mouse E8.5 caudal embryo. Selected progenitor cell marker genes and signalling pathways operating during posterior neural differentiation. B. Basic protocol for in vitro generation and neural differentiation hNMPs (Gouti et al., 2014) and co-expression of Bra/Sox2 proteins on day 3 (NMPs) detected by immunocytochemistry (3 independent experiments). C. RTqPCR assessing relative expression of *Bra* during generation of NMPs (SA121 line). D. RTqPCR for *Pax6* in SA121 cell line cultured as indicated (SA121-line exhibits low-level *Pax6* in hESC, while H9-line does not and so H9 was used for all subsequent experiments). E. RTqPCR for *Pax6* in H9 cells cultured as indicated. F. Schematic of the developed differentiation protocol, including a dual-SMAD inhibition step (dSMADi-RA), and immunocytochemistry for Bra and Sox2 in day 3 NMPs (3 experiments). G. RTqPCR showing *Pax6* expression in H9-line differentiated as in E, +/-100 nM RA from day 3. H. RTqPCR for *Pax6* in cells differentiated as in E, with varying SMAD inhibitor inclusion day 2-4. RTqPCR graphs represent expression normalized to *Gapdh* and relative to hESC levels and constitute 3 independent experiments, error bars are SEMs, here and for all subsequent RTqPCR data.

Most *in vitro* differentiation protocols are informed by our understanding of how the cell type of interest is generated during embryonic development. In the caudal end of amniote embryos, FGF and Wnt signalling act in a positive feedback loop to maintain the elongation of the body axis (Aulehla et al. 2003; Olivera-Martinez and Storey 2007; Wilson et al. 2009). FGF signalling also promotes expression of genes characteristic of CLE including the transcription factor *Nkx1.2* (Delfino-Machin et al. 2005; Sasai et al. 2014). *Nkx1.2* expression extends into the preneural tube (PNT) (Spann et al. 1994; Schubert et al. 1995; Rodrigo-Albors et al. 2016). Here preneural progenitors (PNPs) downregulate *Bra*, transcribe the early neural gene *Sox2*, but as yet lack neurogenic genes such as *Neurog2* and *Pax6* (Scardigli et al. 2001; Scardigli et al. 2003; Bel-Vialar et al. 2007)(Fig 1A). Retinoic acid synthesized in neighbouring paraxial mesoderm mediates the transition from PNPs, repressing expression of *Fgf8* and *Wnts 8a/c* and *3a* (Shum et al. 1999; Diez del Corral et al. 2003; Sirbu and Duester 2006; Olivera-Martinez and Storey 2007; Cunningham et al. 2015) and is then further required for neurogenic gene transcription (Diez del Corral et al. 2003; Ribes et al. 2008)

In addition, to the involvement of these signalling pathways in NMP regulation, inhibition of BMP signalling is required for *Sox2* transcription in the CLE/NSB (Takemoto et al. 2006). In mouse and chick embryos, various BMP and TGFβ antagonists (Noggin, Chordin and Follistatin) are expressed in the anterior primitive streak, emerging notochord and newly formed somites close to posterior neural tissue (Albano et al. 1994; Liem et al. 2000; Chapman et al. 2002). Considered together with the requirement for BMP antagonism for anterior neural induction (Hemmati Brivanlou and Melton 1997; Harland 2000; Kuroda et al. 2004; Linker and Stern 2004) the experiments of Takemoto et al. indicate an on-going requirement for BMP antagonism during the progressive generation of the posterior nervous system.

Almost all in vitro protocols for making NMPs from mouse and human embryonic stem cells (hESCs) involve exposure to various durations of Wnt agonist with or without FGF (Gouti et al. 2014; Tsakiridis et al. 2014; Turner et al. 2014; Lippmann et al. 2015) and one approach has included TGFβ inhibition (to promote loss of self-renewal in human ESC and repress mesendoderm differentiation, after (Chambers et al. 2009)) (Denham et al. 2015). It is well established that efficient induction of anterior neural tissue from hESCs is achieved by exposure to inhibitors of both TGFβ and BMP signalling (known as dual-SMAD inhibition)(Chambers et al. 2009). However, a role for BMP inhibition in the differentiation of neural tissue from NMPs in vitro has not been assessed. Here we show that neural differentiation from human NMPs is promoted by transient dual-SMAD inhibition. We deploy CRISPR-Cas9 engineering to make a reporter for enrichment for human NMPs and provide the first transcriptomic profiling of this cell population and derived spinal cord progenitors.

## Results and Discussion

### Generation of human neuromesodermal progenitors and robust differentiation into posterior neural progenitors by inclusion of transient dual SMAD inhibition

In human ESCs the simplest approach to make NMPs involves removal of self-renewal conditions and exposure to FGF and the Wnt agonist CHIR99021 for 3 days. The NMPs generated in this way were then differentiated into neural progenitors by day 6, following replating and culture in basal media alone (Gouti et al. 2014). We assessed the reproducibility of this protocol (Fig. 1B) to generate *Pax6* expressing neural progenitors. Culturing hESCs in Neurobasal/1x N2/1x B27 medium (N2/B27) supplemented with 20 ng/ml bFgf and 3 µM CHIR99021 for 3 days readily generated Sox2/Bra co-expressing NMP cells (Fig. 1B, C). However, subsequent differentiation after cell dissociation and replating in just N2B27 at the end of day 3 (D3), did not generate *Pax6* positive cells by end of day 6 (D6) (assessed in two hESC lines, SA121 and H9) (Figs 1D, E). We therefore next included all-trans retinoic acid (RA) 100 nM in this differentiation protocol from the beginning of day 4 (D4). However, this regime also did not elicit reproducible expression of *Pax6* by D6 (Fig. 1D, E). This contrasted with a positive control for *Pax6* transcription provided by a protocol for inducing anterior neural progenitors, exposure to Noggin 50 ng/ml and the TGFβ receptor type 1 inhibitor SB431542 10 µM following removal of self-renewal conditions (dual SMAD inhibition, (Chambers et al. 2009), Fig. 1D).

This inability to induce *Pax6* from NMPs might reflect inherent differences between hESC lines (MasterSheff in Gouti et al 2014, compared with H9 this study), but could also involve variant culture conditions. In particular, while we used EDTA and gentle mechanical agitation to replate at D3, these researchers used Accutase, which can damage extra-cellular matrix and membrane proteins (Beers et al. 2012). This might then reduce cell-cell signalling and could mimic inhibition of BMP signalling, as reported on dissociation of Xenopus animal cap ectoderm (Wilson and Hemmati-Brivanlou 1995). We therefore next employed dual SMAD inhibitors from the beginning of D3 to the end of D4. This period aimed to mimic exposure to endogenous TGFβ inhibitors experienced by cells in the CLE and PNT in the amniote embryo (Fig. 1A). Inclusion of this step did not alter induction of NMPs on D3, as judged by generation of Sox2/Bra co-expressing cells (Fig. 1F and see flow cytometry data Fig. S1), but, this step alone was still not sufficient to promote reproducible *Pax6* expression by D6 (Fig 1G). However, in combination with subsequent exposure to RA from D4, robust *Pax6* was induced in this timeframe (Fig. 1G). Importantly, inclusion of either Noggin or SB431542 alone with RA was not effective (Fig. 1H), indicating that dual SMAD inhibition is required to augment neural differentiation in this context. The reproducibility of this dual SMAD inhibition and RA protocol (dSMADi-RA) (Fig. 1F) was further demonstrated by rapid induction of *Pax6* in a hiPSC line (Fig S2: ChiPS4).

To characterize this dSMADi-RA differentiation protocol we analyzed the expression dynamics of key cell state marker genes using quantitative reverse transcription PCR (RT-qPCR). Pluripotency genes *Nanog* and *Oct4* were dramatically reduced from hESC to D3(NMP) and transcripts were lost quickly as these cells differentiated (Fig. 2A), as observed in mouse and chick embryo and mouse ESC-derived NMPs (Tsakiridis et al., 2014, Gouti et al., 2014). D3(NMPs) were characterized by high-level *Bra* and *Cdx2* transcription (Fig 2B). As in mouse ES cell-derived NMPs, *Sox2* transcripts were lower in D3(NMPs) than in hESCs, despite high-level Sox2 protein in NMPs (Gouti et al 2014; Turner et al 2014) (Fig. 1F, 2B). *Cdx* genes regulate signalling that maintains the mouse NMP cell state and also induce expression of posterior Hox genes, which confer anterior-posterior identity (Young and Deschamps 2009; Young et al. 2009; Gouti et al. 2017). NMPs and their derivatives generated with this protocol expressed *Hoxb4* and *Hoxc6*, indicative of thoracic identity (Fig. 2C). Although expression of these genes commenced in D3(NMPs), their transcription increased during neural differentiation and so must also be indicative of the identity of neural progenitors generated in these conditions. In the embryo, differentiation from NMPs to neural progenitors involves downregulation of *Bra* and entry into a transitional preneural cell state (Fig. 1A), which is detected in vitro by persisting expression of *Wnt8a/c* and *Nkx1.2* (Fig. 2E). As these genes decline *Pax6* is then transcribed, rising to a peak at D8 (Fig. 2D). This suggests that neural progenitors arise between D5 and D8. This protocol therefore provides an assay in which to investigate the human NMP cell state and how spinal cord progenitors are formed.

**Figure 2.**
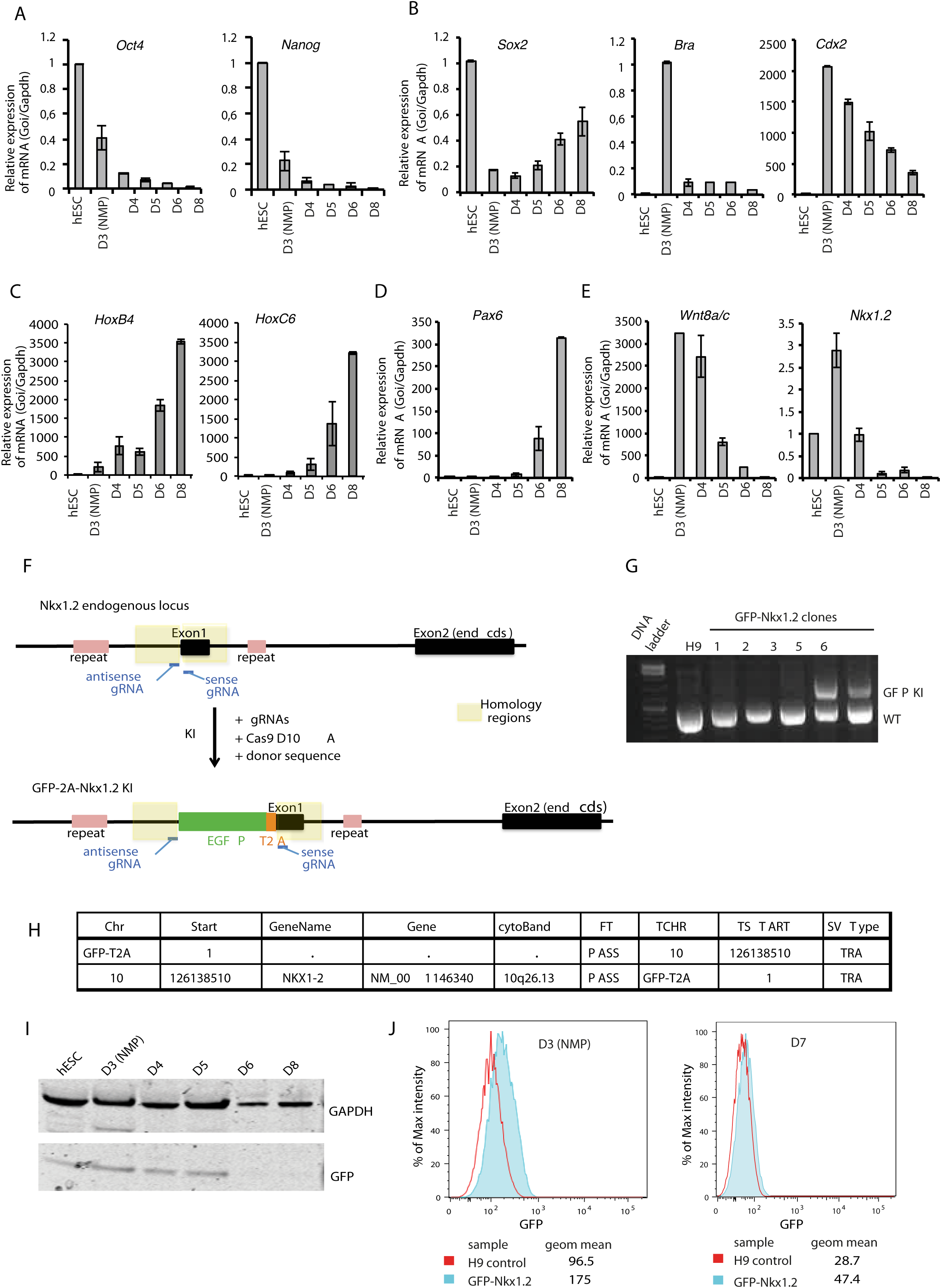
RTqPCR for selected genes during dSMADi-RA differentiation and generation of a GFP-Nkx1.2 reporter line. A-E. RTqPCR assessing relative expression of key marker genes in H9-cells exposed to dSMADi-RA protocol (Fig. 1F). A. Declining expression of pluripotency genes *Oct4* and *Nanog*. B. *Sox2*, *Bra* and *Cdx2* expression dynamics. C. *HoxB4* and *HoxC6* during differentiation. D. Expression of neurogenic neural progenitor marker *Pax6*. E. *Wnt8a*/*c* and *Nkx1*.*2*, characteristic of preneural progenitors and NMPs. F. Experimental strategy schematic: H9 hESC were engineered using CRISPR/Cas9, knocking-in the GFP-T2A sequence upstream exon 1 of *Nkx1.2*. Positions of the gRNAs, and homologous regions used in the repair template are indicated. G. PCR amplification of the *Nkx1.2* locus using primers framing the insertion site. H9: untransfected control, 1-3: GFP negative clones, 5 and 6: clones containing the GFP insertion (GFP KI = knockin, WT = wildtype allele). H. Whole genome sequencing of GFPNkx1.2 clone 5. Structural variation analysis relative to GFP-T2A sequence: FT: per sample genotype filter; TCHR: chromosome for the translocation breakpoint coordinate; TSTART: translocation breakpoint coordinate; SVType: structural variation type; TRA: translocation. I. Western blot of GFP during differentiation of the GFP-Nkx1.2 line. J. Flow cytometry of GFP expression at day 3 and day 7: % of maximum-intensity for GFP-channel is plotted, representative values of at least 2 experiments.

### Generation of a human Nkx1.2 reporter cell line

Cell populations generated in vitro are inevitably heterogeneous and so we next made a reporter line that could be used to enrich for NMPs. We took advantage of CRISPR/Cas9 technology (Komor et al. 2017) to engineer H9 hESCs to express GFP under the control of the endogenous *Nkx1.2* promoter. This homeo-domain containing transcription factor is highly expressed in NMPs (CLE and NSB) in the embryo and persists in preneural cells (Fig. 1A, 2E) but is not detected in non-neural tissues (Spann et al. 1994; Schubert et al. 1995; Rodrigo-Albors et al. 2016). We reasoned that selection for high *Nkx1.2* expression on D3 when *Bra* transcripts are high would enrich for NMPs. To this end a GFP-T2A sequence (Kim et al. 2011) was knocked-in to the *Nkx1.2* locus in-frame just upstream of exon1 (Fig 2F and see Materials and Methods). Correct targeting was confirmed by PCR across the integration site and subsequent fragment sequencing (Figs 2G, S3). Whole genome sequencing and structural variation analysis of this data further confirmed that the *Nkx1.2* gene was the only locus modified by integration of GFP-T2A (Fig. 2H). Using CRISPR-Cas9 approach we thus generated a GFP-Nkx1.2 hESC line bearing a mono-allelic insertion of the GFP-T2A specifically in the *Nkx1.2* locus.

Differentiation of this GFP-Nkx1.2 reporter line using the dSMADi-RA protocol was then characterized by Western blot; revealing GFP expression up to day 5 (Fig 2I), including low-level GFP in hESC, (consistent with detection of *Nkx1.2* in H9 hESCs) (Fig. 2E). Flow cytometry (without GFP antibody) further confirmed GFP expression at D3 in GFP-Nkx1.2 cells compared with auto-fluorescence profile of wild-type H9 differentiated in parallel, which was then lost as cells differentiate (D7) (Fig. 2J). To confirm that *Nkx1.2* locus modification did not impair differentiation we used immunocytochemistry and flow cytometry to assess Sox2/Bra co-expression on D3 (Fig. S1) and RT-qPCR (Fig. S4) to profile expression of marker genes during dSMADi-RA differentiation. These analyses indicated that the engineered line made NMPs and that its differentiation was comparable to that of the parental H9 line (Figs S1, S4, 2A-E). Similar results were obtained with a second GFP-Nkx1.2 line, demonstrating the reproducibility of this approach (Fig. S5).

### Identity and conservation of human NMP transcriptional signature

We next used this GFP-Nkx1.2 cell line to select for high GFP-expressing cells on D3 using FACS (see Materials and Methods) and generated RNA-Seq data for D3 and D8 cell populations. This was also compared with published RNA-Seq data for H9 hESCs (Chu et al. 2016). Genes more highly expressed on D3 were identified by comparison with hESCs and D8 neural progenitors (Fig. 3A, Table S1). This included expected NMP-associated genes *Bra, Cdx1, Sp5, Wnt8A/C, Fgf17,* but also new genes, such as membrane protein of unknown function *Unc93a* and *GPRC5a*, an orphan G-protein-coupled receptor responsive to retinoid signalling (Cheng and Lotan 1998). Some enriched genes (*Fgf17, GPRC5A, Unc93A*) were then validated by RT-qPCR, including a gene not in the top list (*Shisha3*), which attenuates FGF and Wnt signalling (Yamamoto et al. 2005) (Fig. 3B).

**Figure 3.**
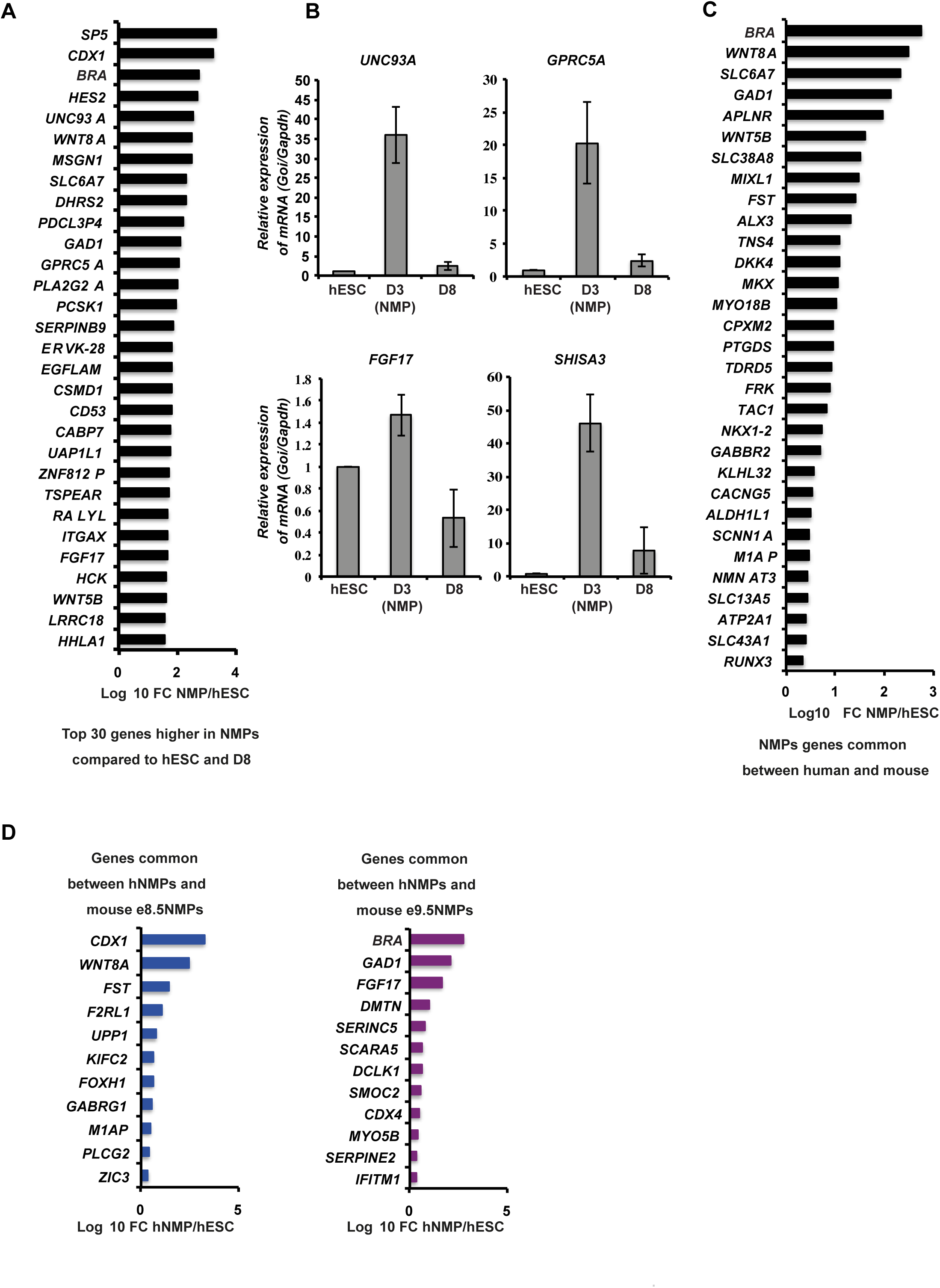
Characterization and conservation of Human D3(NMP) transcriptome. A. Genes preferentially expressed in human D3(NMPs) compared to hESC and hD8 neural progenitors. Genes were considered preferentially expressed in hD3(NMPs) when >2-fold-change between hD3(NMPs) and both hESC and hD8 (full list TableS1). B. RTqPCR for subset of D3(NMP)-enriched genes. C. Comparison of enriched genes identified in human (this study) and bulk-RNA-Seq of mESC-derived NMPs (Gouti et al., 2014). D. Comparison of NMP-enriched genes in human (this study) with mouse embryo eNMP transcriptional signatures obtained by comparing scRNA-Seq data for e8.5 and e9.5 embryos (Gouti et al., 2017).

This human D3 gene list compared was next with that for genes uniquely upregulated in *in-vitro*-derived mouse NMPs (Gouti et al. 2014). This identified 31 conserved NMP genes (Fig. 3C). These include transcription factors known to be expressed in NMPs*, Bra, Nkx1.2* and *Mixl1*, but also newly implicate *Mkx* (mohawk /Irx1L), (Liu et al. 2006), *Alx3* (Beverdam and Meijlink 2001) and *Runx3* as transcriptional regulators of this cell population. Predicted signalling pathways, Wnt (*Wnt8A, Wnt5A, Dkk4*) and TGFβ antagonism (*Fst, Follistatin*) were also represented, along with genes implicating new signalling activities. These include 4 solute carriers mediating citrate (SLC13A5) or amino acid transport (SLC38A8, SLC43A1, SLC6A7). *SLC6A7* is a member of the gamma-aminobutyric acid (GABA) neurotransmitter gene family and strikingly two further genes mediating GABAergic signalling are also conserved: *GAD1* (glutamic acid decarboxylase), which synthesizes GABA from glutamate and is transcribed in the mouse tailbud (Maddox and Condie 2001) and GABA receptor *GABBR2/*GPRC3B. In neurons, GABA-B receptors can trigger inactivation of voltage-gated calcium channels (Padgett and Slesinger 2010). Two further conserved NMP genes, *CACNA1C* (a calcium-channel auxiliary subunit/CaV1.2) implicated in maintaining calcium-channel inactivation (Soldatov et al. 1997) and *ATP2A1* (a calcium transporting ATPase) which maintains low cytoplasmic calcium (Shull et al. 2003), may be additionally operating via different mechanisms to attenuate intra-cellular calcium levels. This is consistent with the requirement for calcium signalling for neural induction identified in chick embryos (Papanayotou et al. 2013) as indicated by *Sox2* transcription, which is indeed detected at low-levels in NMPs and then rises in neural progenitors.

As there are not only species differences between these data sets, but also in vitro protocol variation, we additionally compared the human D3(NMP) molecular signature with those obtained for mouse embryonic NMPs at E8.5 and E9.5 using single-cell-RNA-Seq (Gouti et al. 2017). This identified 23 genes conserved between mouse embryo and *in vitro* derived human NMPs (Fig. 3D). This again included *GAD1* and another GABA receptor, *GABRG1*, belonging to the type-A family, shown to regulate stem cell proliferation (Andang et al. 2008). GABA biosynthesis is an output of the tricarboxylic acid (TCA) cycle, input to which can come from glycolytic metabolism, which has recently been shown to operate in tailbud progenitor cell populations (Bulusu et al. 2017; Oginuma et al. 2017). It will therefore be important in the future to understand the relationship between this metabolic state and GABA production in NMPs (Fig. 3D).

### Transcriptomic characterization of the differentiation protocol

These RNA-Seq data also helped to characterize cell types generated with the dSMADi-RA differentiation protocol. The mesendoderm marker *Sox17* was not detected, nor were transcripts from anterior neural genes (*Foxg1, En2* and *Dlx2)* in any condition (<10 reads), while *Otx2*, which is initially expressed in the early epiblast and the primitive streak (Ang et al. 1996; Henrique et al. 2015), declines sharply from hESCs (Fig. 4A). This is not surprising given hESC exposure to FGF and Wnt signalling for 3 days to generate NMPs, at which time cells begin to express a range of Hox genes, including A1, B1, B4 and A7, (Fig. 4B). In this assay, therefore, NMPs possess a posterior identity prior to their direction along the neural differentiation pathway. Components of signalling pathways regulating NMPs exhibited expected gene expression profiles (Figs 4C, D, E). High-level transcription of neural progenitor and neurogenic genes (Fig 4F) was detected on D8 and correlated with increased retinoid signalling reported by *RARb* transcription (Fig. 4G). The expression of both BMP and Shh pathway genes (Fig. 4H,I) on D8 suggested that induced spinal cord progenitors are exposed to dorsal (BMP) and ventral (Shh) patterning signals. However, while dorsal neural progenitor and neural crest associated genes were expressed along with some more ventral progenitor genes (Fig. 4J), ventral-most markers *Nkx2.2* and floor plate marker *FoxA2* were not detected at D8 (data not shown). The early transcription of neural crest genes in this differentiation assay further suggests that as in the elongated embryonic body axis and in mouse ES-derived *in vitro* spinal cord assays, dorsal progenitor cell types emerge prior to ventral progenitors (Meinhardt et al. 2014).

**Figure 4.**
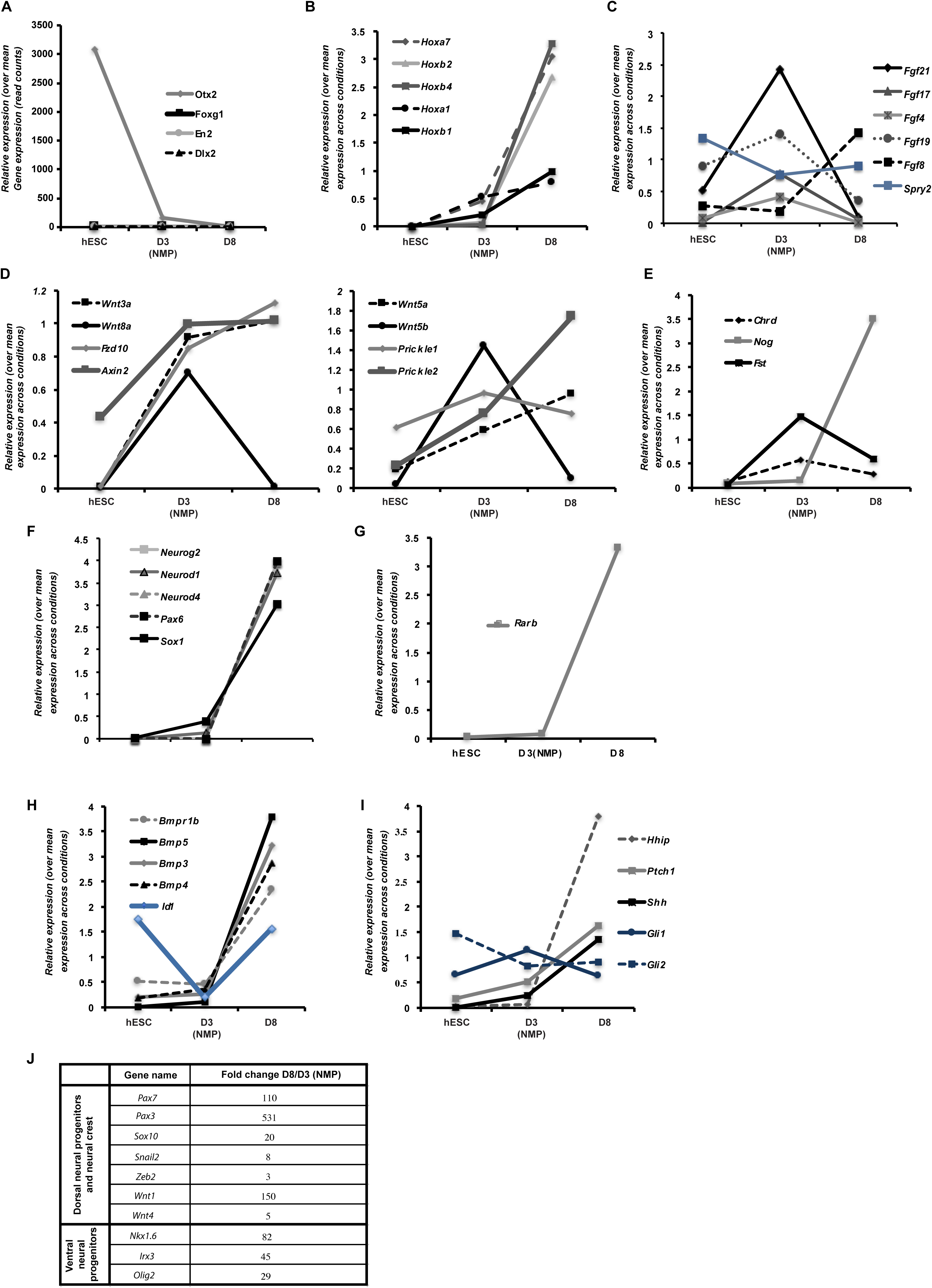
Expression of selected genes across 3 conditions analysed by RNA-Seq. A. Anterior neural marker genes, presented as read counts. B. Main *Hox* genes expressed at NMP stage. Selected components of C. FGF, D. Wnt signaling pathways. E. Selected BMP/TGFβ inhibitors. F. Neural progenitor and neurogenic genes and G. Retinoid receptor *RAR*b during NMP differentiation. Selected components of H. BMP and I. Shh signalling pathways. B-I relative expression of each gene normalized to its mean expression across all conditions J. Table of neural crest, dorsal and ventral progenitor genes induced during dSMADi-RA differentiation. Fold change between D8 and D3(NMP) time points is shown. SEMs for each gene shown in Figs S6A, S6B, Table S2.

Guided by signalling in model vertebrate embryos we have devised a protocol for the robust differentiation of human spinal cord progenitors from neuromesodermal progenitors, which can be used for future mechanistic and translational approaches, including development of human neuroepithelial cell behaviour assays. The GFP-Nkx1.2 reporter line ensured selection of high *Nkx1.2* expressing cells on D3 and has the potential to be further engineered to report for *Bra*, select for later Nkx1.2+/Bra-cells and so identify earliest changes in progression towards neural differentiation. These RNA-Seq data not only served to validate this differentiation protocol and uncover a conserved NMP transcriptional signature, but also identified potential new regulatory mechanisms, including GABA and calcium signalling in the maintenance of the NMP cell state.

## Materials and Methods

### Human ES cell culture and differentiation

Human ES cells (H9, sa181, sa121) and human iPS cells (ChIPS4) were maintained as feeder-free cultures in DEF-based medium (Cellartis DEF-CS) supplemented with bFGF (30 ng/mL, Peprotech) and Noggin (10 ng/ml, Peprotech) on fibronectin coated plates, and enzymatically passaged using TryLEselect (thermofisher). For passaging, the medium was supplemented by addition of the Rho kinase inhibitor Y-27632 (10 mM, Tocris). For differentiation assays, PSC were plated on Geltrex matrix (20µg/cm^2^, LifeTechnologies) at a density of 40000 cells/cm^2^ and shifted to N2B27 medium after 24h. Cells are differentiated for 3 days in N2B27 supplemented by 3 µM Chiron99021 (Tocris) and 20 ng/ml bFgf (PeproTech) to obtain NMPs. For further differentiation, NMPs were passaged using PBS-EDTA 0.5 mM and seeded back at controlled density (200000 cells cm^2^) on Geltrex (20µgcm^2^, LifeTechnologies) in the presence of Y-27632 (10 mM, Tocris). From passaging, cells are then cultured in N2B27 containing 100 nM RA for the indicated time to obtain later stage progenitors. In addition to the above, 50 ng/mL Noggin and 10µM SB431542 are added from day 2 to day 4. All experiments with hESCs were approved by the UK Stem Cell Bank steering committee (SCSC14-28 and SCSC14-29).

### RTqPCR

Total RNA was extracted using the RNEasy mini kit (Qiagen), following the manufacturer’s instructions, with the addition of a DNAse digestion step performed on the column for 15 min with RQ1-DNase (Promega). After initial denaturation for 5 min at 70°C in presence of 1 µg random primers, 500 ng of RNA per sample were reverse transcribed for 1h in 20 µL reaction volume containing 0.5 mM dNTPs, 5 mM MgCl2, 1X ImProm-II RT buffer, 20 U RNasin and 160 U of ImProm-II RT (Promega). Samples were incubated for 15 min at 70°C to stop the reaction. qPCR analysis was performed using primers described in table 6 on either a Mastercycler RealPlex2 (Eppendorf) or an AriaMX (Agilent) device in presence of PerfeCTa SYBR Green SuperMix for iQ (Quanta Biosciences) or BrilliantIII SYBRgreen PCR MasterMix (Agilent) respectively. Relative expression was calculated using the ΔΔCt method, normalizing each gene of interest to *Gapdh* levels.

### Western blot

Western blots were performed using standard protocols. Briefly, proteins were extracted using RIPA buffer (150 mM sodium chloride, 1.0% Triton X-100, 0.5% sodium deoxycholate, 0.1% SDS (sodium dodecyl sulphate) and 50 mM Tris, pH 8.0). Cell extract was incubated on ice for 30 min in presence of DNAse (Universal Nuclease, Pierce) and spun down for 20 min at full speed. Protein concentration in supernatant was determined using a Bradford Assay with a BSA standard curve ranging from 0-2 mg/ml. The samples were diluted in NuPage 4x sample buffer (LifeTechnologies) and loaded onto a 4-12% gradient gel (Novex NuPAGE, LifeTechnologies). Western blots were performed using standard procedures and antibodies used at the following concentrations: anti-GAPDH 1 µg/mL (ab9484, abcam), anti-GFP 1 µg/mL (ab6673, abcam). Detection was performed with anti-goat Dylight 6800 conjugate (1:10000, LifeTechnologies) and anti-mouse DyLight™ 800 conjugate (1:10000, LifeTechnologies) on a LI-COR imaging device (BioSciences).

### Immunofluorescence microscopy

Cells were fixed by adding formaldehyde to a final concentration of 3.7% in PBS, then permeabilized and blocked in PBS/0.1% TritonX-100/4%(w/v)BSA. Incubation was performed at 4°C overnight with primary antibodies at the following concentrations: goat anti-Brachyury 1 µg/mL (AF2085, R&D), rabbit anti-Sox2 5 µg/mL (ab5603, Millipore). Fluorochrome-conjugated secondary antibodies used were the following: anti-goat Alexa647-conjugated 4 µg/mL (A21447, Invitrogen), anti-rabbit Alexa488-conjugated 4 µg/mL (A21206, MolecularProbes). Observations were carried out with a DeltaVision fluorescence microscope (GE Healthcare), and images were acquired using the softWoRx software.

### Flow cytometry analysis of protein expression profile

Cells were harvested using TryLEselect, fixed for 10 min in 4% paraformaldehyde and re-suspended as single cells in PBS 1%BSA. An additional 10 min methanol fixation step was added for Sox2 and Brachyury detection. Primary antibodies were incubated for 1 h at room temperature in PBS 4% BSA, cells were then washed and incubation with secondary antibodies performed for 30 min at room temperature. Antibody used are the followings: goat anti-Brachyury 1 µg/mL (AF2085, R&D), rabbit anti-Sox2 5 µg/mL (ab5603, Millipore), anti-goat Alexa647-conjugated 2 µg/mL (A21447, Invitrogen), anti-rabbit Alexa488-conjugated 2 µg/mL (A21206, MolecularProbes). After washes, fluorescence was measured on a FACSCanto cytometer (BD Bioscience) and results analyzed using FlowJo software. Quadrant gates used to estimate the percentage of positive cells were designed based on fluorescence levels detected in the control samples processed without primary antibodies.

### GFP-Nkx1.2 engineering

The donor plasmid construct, pDonorNkx1.2NterKI, was synthesized by GeneArt. The vector is based on a pMK-RQ backbone and contains a Kanamycin resistance cassette and the GFP-T2A insert flanked by 500 bp homology arms for recombination to the Nkx1.2 5’ end. The second plasmid used, px335Nkx1.2NterKIas, encoded the Cas9D10A nickase (Cong et al., 2013) and the antisense gRNA (asgRNA GCCCACGGGCCGGCGGTCGG). A third plasmid, pBABEDpU6Nkx1.2NterKIs, included the sense gRNA (sgRNA GCTGGCATGGCAGGACGGCG) and a puromycin resistance cassette to select transfected cells. CRISPR-Cas9 mediated gene targeting was performed as follows. H9 hESC were dispersed to single cells using TryLEselect (thermofisher) and re-suspended in DEF medium in the presence of Y-27632 (10mM, Tocris). For transfection, 5x10^6^ cells were pelleted by centrifugation at 300 xg for 3 minutes, washed with PBS and re-suspended in 100µL buffer R (Neon Transfection Kit, Thermofisher). 4ng of pDonorNkx1.2NterKI, 2ng of px335Nkx1.2NterKIas and 2ng of pBABEDpU6Nkx1.2NterKIs was added to the mix. Electroporation was performed with the Neon Transfection System (Thermofisher) using the following parameters: 1 pulse, 1150V, 30ms. Transfected cells were plated and allowed to recover for 36 hours, puromycin selection was applied for a further 36 hours. Clones were left to grow until easily visible, hand-picked and seeded back in 96 well plates before being amplified. Screening of the clones for GFP integration was performed by PCR using primers amplifying across the insertion sites (GFPcheckFw1+GFPcheckRev1, see the table below for sequences) (Fig. 2G). Correctly targeted integration of the GFP-T2A sequence was checked in 40 transformed hESC clones by PCR across the integration site. Overlapping PCR amplification products spanning the locus from outside the homologous region to inside the GFP sequence on both sides of the integration were sequenced (GFPcheckFw1+GFPcheckRev2 and GFPcheckFw2+GFPcheckRev1, see the table below for sequences). Five clones were found to include the GFP-T2A sequence at the correct locus and these were all heterozygous for GFP-Nkx1.2 (Fig. 2G). PCR bands obtained for GFP-Nkx1.2 clone 5 were sequenced to check for integrity of the recombination borders and absence of mutations. Results were combined and detailed sequence of the engineered allele was obtained (Fig. S3).

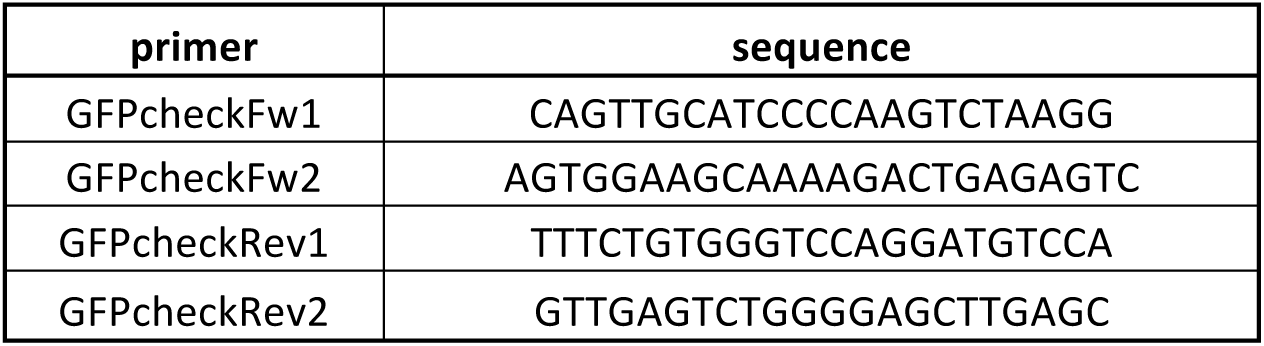

### Whole Genome Sequencing

gDNA was extracted from GFP-Nkx1.2 hES cells using the DNeasy Blood and Tissue Kit (Qiagen), according to the manufacturer’s instruction. Whole Genome Sequencing was performed by Novogene. Briefly, a library was generated from 1 µg gDNA using Truseq Nano DNA HT sample preparation Kit (Illumina) following manufacturer's recommendations and sequenced on an Illumina platform. After quality control, BWA (version 0.7.8‐r455) was used to align reads to the genome, using the 1000Genomes (GRCh37+decoy) human as reference. BAM files were sorted using SAMtools (version 1.0) and read duplicates identified using Picard (version 1.111). Structural variation (SV) analysis was done using Delly (v0.7.2) (Rausch et al. 2012), and ANNOVAR (version 2015Mar22) was used to annotate the SV. An average coverage of 33X was obtained (depth >20X for 92% of bases).

### Cell purification for RNA-Seq analysis by FACS

Cells were sorted on a BD Influx (Becton Dickinson) cell sorter using the 100 µm nozzle. FSC vs SSC was used to identify live cells and then FSC-A vs FSC-W to identify single cells. The GFP positive cells were identified using 488 nm laser light and the parameters GFP (530/40) and PE (580/30). The gate to identify GFP positive cells was set using a GFP negative control (H9 cells differentiated in parallel) and events that fell into this gate were sorted to more than 97% purity. 1.5 million GFP positive cells sorted at day 3 were used per sample for RNA extraction.

### Library preparation for RNA-Seq and Sequencing

Total RNA was extracted using the RNEasy mini kit (Qiagen), following the manufacturer’s instructions, with the addition of a DNAse digestion step performed on the column for 15 min with RQ1-DNase (Promega). RNA concentration was measured on a Qubit device using Qubit RNA BR assay kit (ThermoFisher), and quality was checked on a TapeStation instrument (Agilent). Individually labeled libraries were prepared from 1 µg of RNA per sample using the TruSeq^^®^^ Stranded mRNA Library prep kit (Illumina), according to the manufacturer’s instruction. 2 µL of Spike-ins (1/100 dilution ERCC Spike-in controls Mix1) was added per sample. Libraries were pooled and sequencing was performed on a NextSeq (Illumina) at the Tayside Center for Genomic analysis (Ninewells, Dundee) as follows: high output run, 2x75bp paired end sequencing, between 35 and 46 million uniquely mapped reads obtained per sample (12 samples multiplexed). RNA-seq data are available in the Array express database (http://www.ebi.ac.uk/arrayexpress) under accession number XXX

### RNA-Seq analysis

RNA-Seq reads were mapped to the reference genome (version GRCh38, release 87) using STAR 2.5.2b, using stranded option. Typically, about 92% of reads were mapped uniquely (except for D3(NMP) replicate 4, where uniquely mapped reads were at 86.8%). Read counts per gene were found in the same STAR run. Data from (Chu et al. 2016) were re-analysed in the same fashion, however, we point out that these were single-end non-stranded reads. For the following analysis, 4 biological replicates were used for D3(NMP) and 2 for D8 samples. Differential expression was performed with edgeR 3.16.5, for each pair of conditions independently. Benjamini-Hochberg multiple-test correction was applied to test p-values. Human NMP genes (Fig. 3A) were determined by selecting genes using the following criteria: at least 10 read counts in D3(NMP), significantly enriched (p value < 0.01) in D3(NMP) compared to both hESC and hD8 samples, with a foldchange >2. Time-dependent properties of genes were studied using intensity profiles hESC-D3(NMP)-D8. Each point in the profile is a DESeq-normalized mean gene count across replicates. To make profiles comparable, they were normalized to their mean across conditions, so the mean of each normalized profile is 1.

## Acknowledgements

We thank members of the Storey Laboratory and Professor Carol MacKintosh for critical reading of this manuscript. This research was supported by a Wellcome Trust Investigator Award to KGS (WT102817AIA).

---

**Figure S1.**
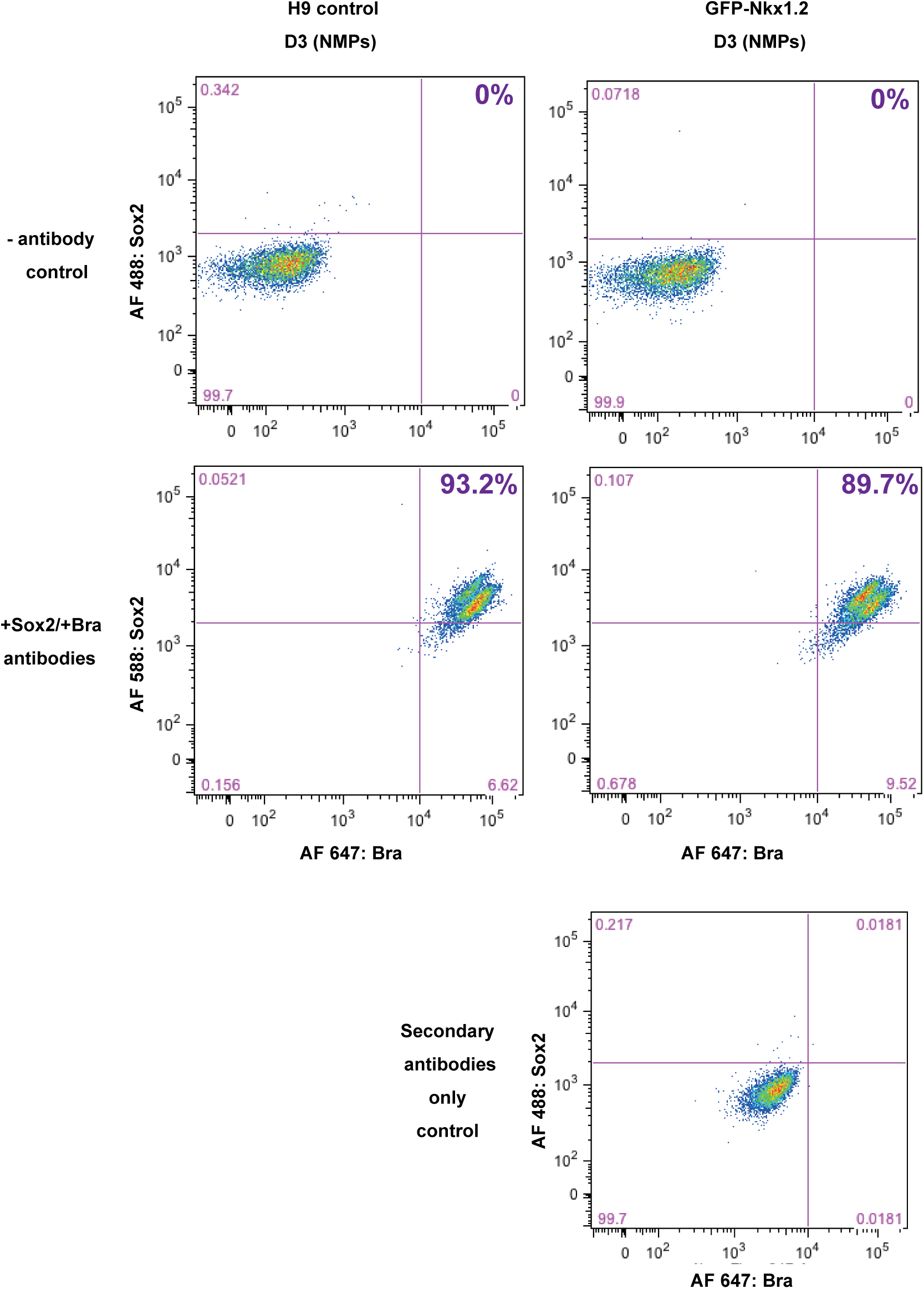
Co-expression of Sox2 and Bra proteins in hD3(NMPs) Expression of Sox2 and Bra was analyzed by flow cytometry in D3(NMPs) derived from H9 cell line (left panels) and H9-GFP-Nkx1.2 cell line (right panels). Upper panels: no antibodies control, middle panels: staining with anti-Sox2 and anti-Bra antibodies, bottom graph: secondary antibodies alone (control). The quadrant for quantification of co-expression levels was defined based on fluorescence observed without primary antibody application (bottom graph). The percentage of co-expression for each panel is indicated at the upper right corner (purple). Representative experiment of 2 independent experiments.

**Figure S2.**
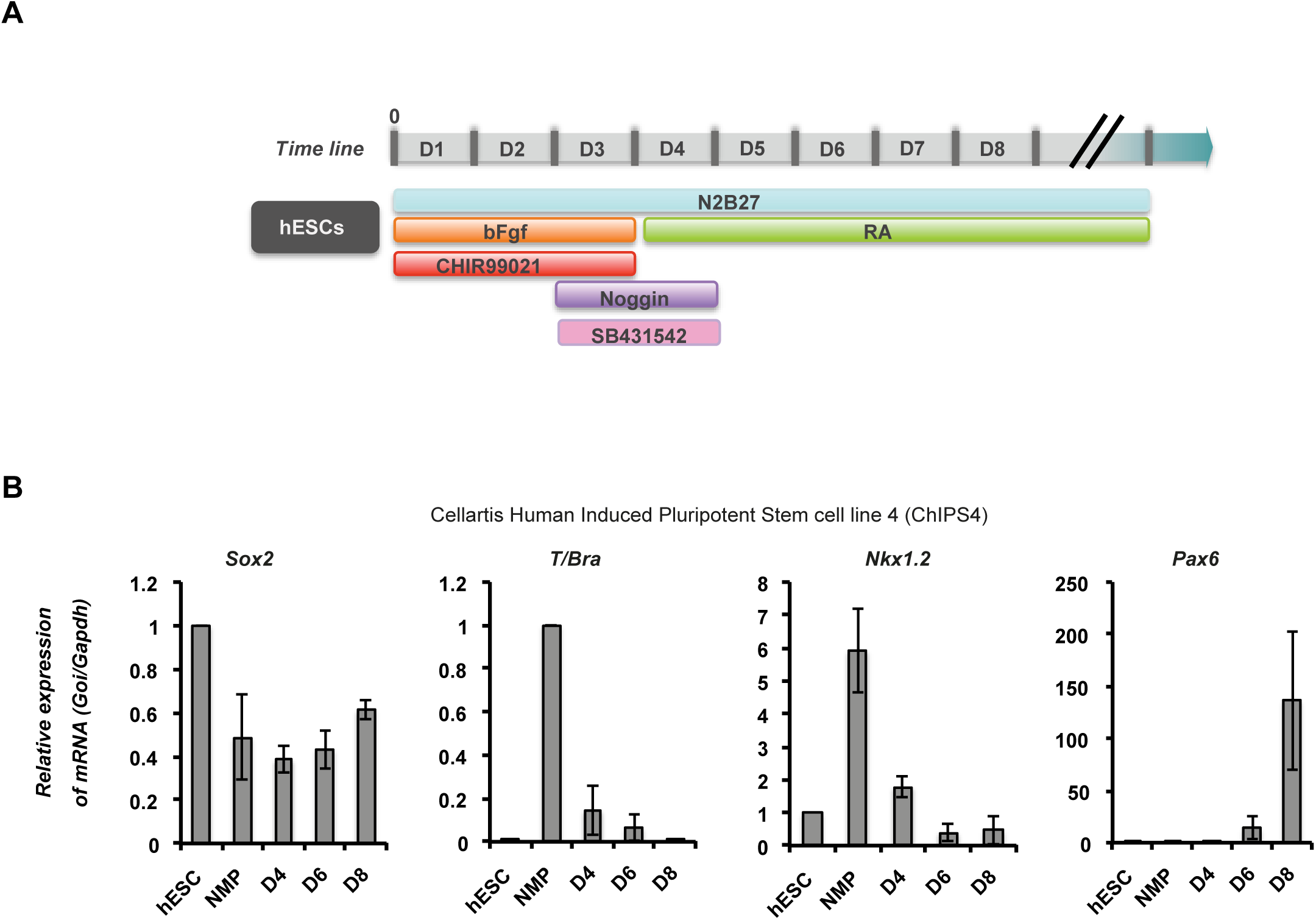
Robust differentiation of posterior neural progenitors from neuromesodermal progenitors in an iPS cell line. A. Schematic representation of the differentiation protocol used on the ChiPS4 cell line, including a dSMADi step from end of day 2 to end of day 4. B. Expression profile of selected genes over time in chiPS4 cells submitted to the differentiation protocol presented in A. Graphs represent the expression of each individual gene normalized to *Gapdh* and relative to hiPSC levels. Average of 3 independent RTqPCR experiments, error bars: standard deviation.

**Figure S3.**
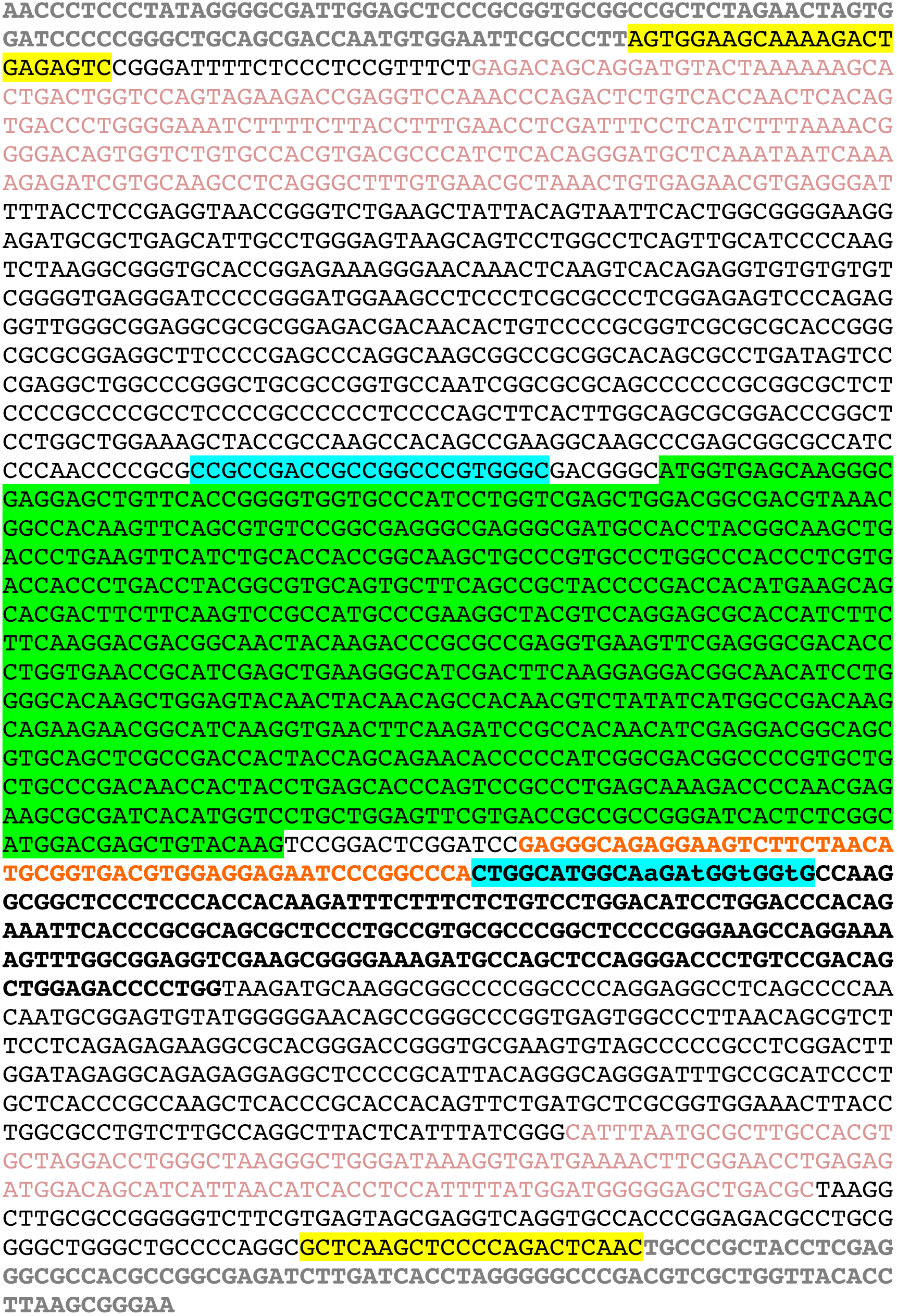
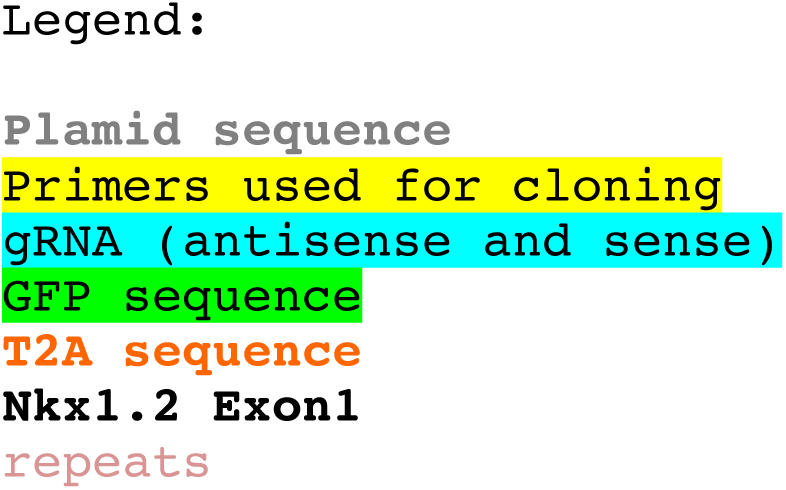
Sequencing of the GFP-T2A insertion site in the correctly targeted clone used in this study. Grey: plasmid sequence, highlighted yellow: Primers used for cloning, highlighted blue: gRNA (antisense and sense), highlighted green: GFP sequence, orange: T2A sequence, bold black: Nkx1.2 Exon1, peach: repetitive sequences.

**Figure S4.**
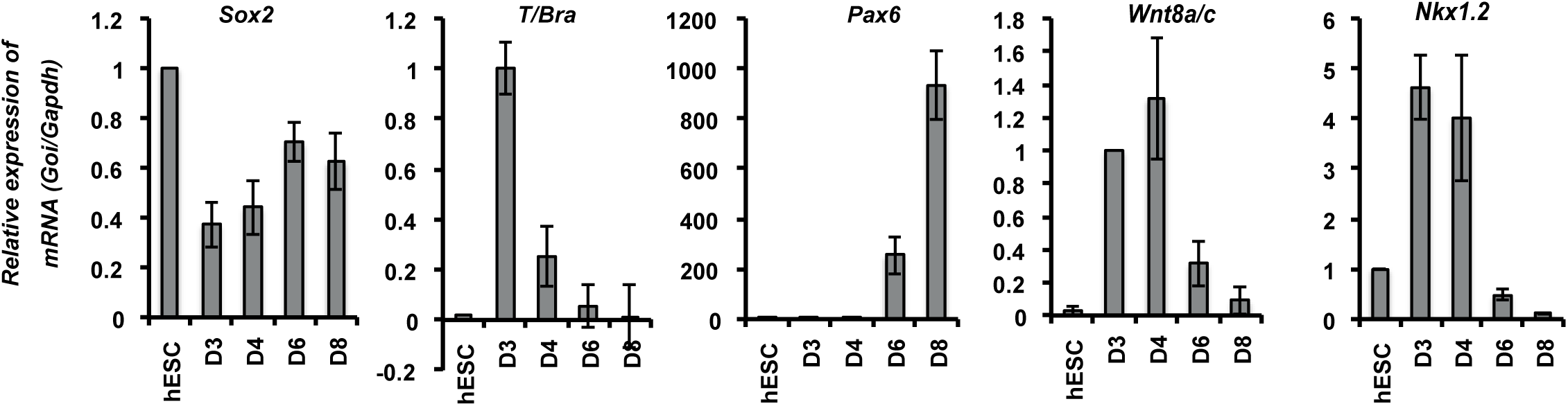
Diffentiation evaluation of the GFP-Nkx1.2 clone used in this study. Expression of selected marker genes was analyzed by RTqPCR during differentiation of the GFPNkx1.2 line following the protocol presented Figure 1F. Graphs represent the expression of each individual gene normalized to *Gapdh* and relative to hESC levels. Average of 3 independent RTqPCR experiments, error bars: standard deviation.

**Figure S5.**
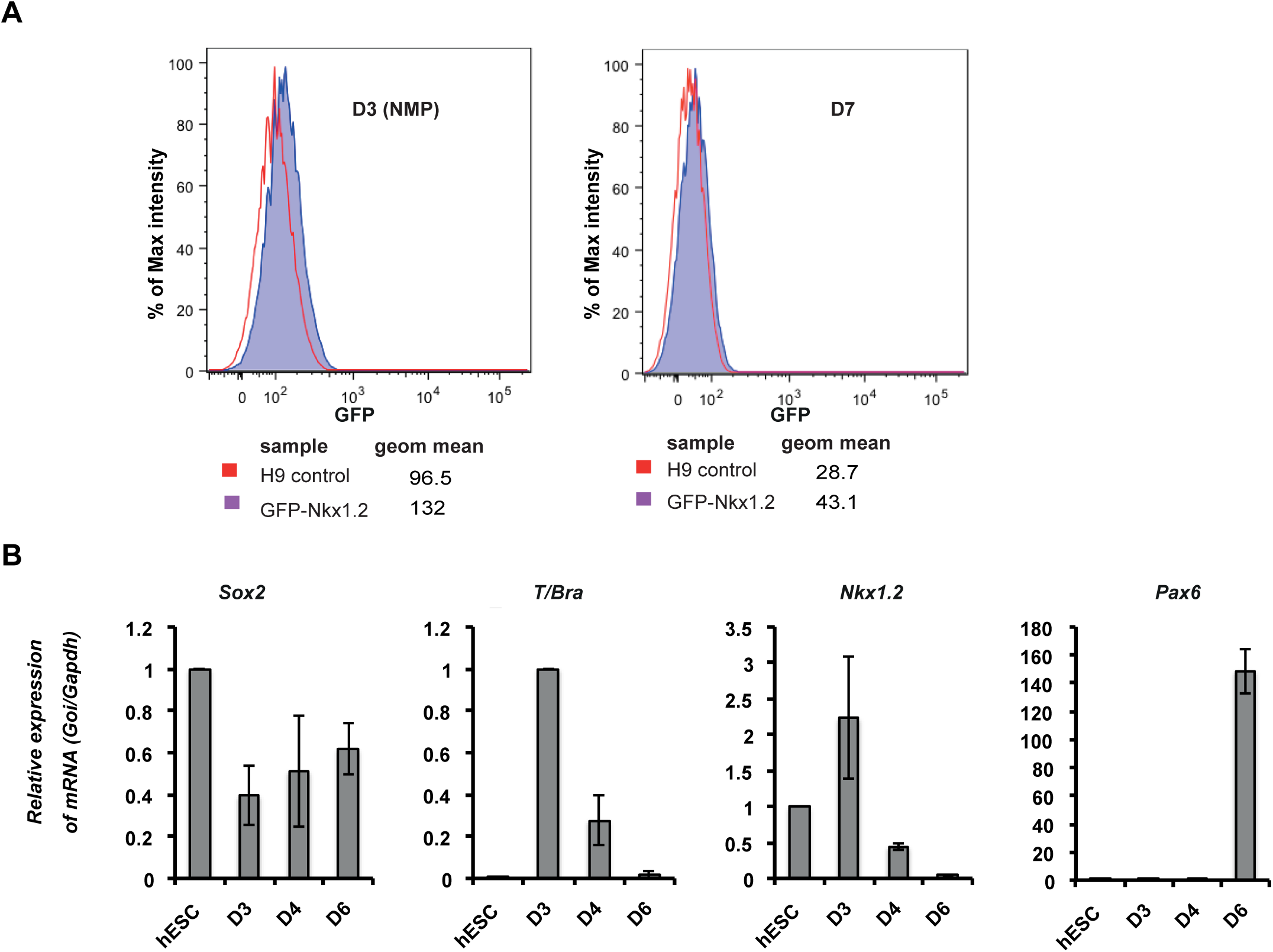
GFP expression and differentiation profile of a second properly targeted GFP-Nkx1.2 clone. A. Flow cytometry analysis of GFP expression at day 3 (NMP) and day 7 of the differentiation protocol. % of maximum intensity for GFP channel is plotted, geometric mean for each peak is indicated. B. Expression of selected marker genes analyzed by RTqPCR during differentiation of the second GFP-Nkx1.2 clone following the protocol presented Figure 1F. Graphs represent the expression of each individual gene normalized to *Gapdh* and relative to hESC levels. Average of 3 independent RTqPCR experiments, error bars: standard deviation.

**Figures S6A and S6B.**
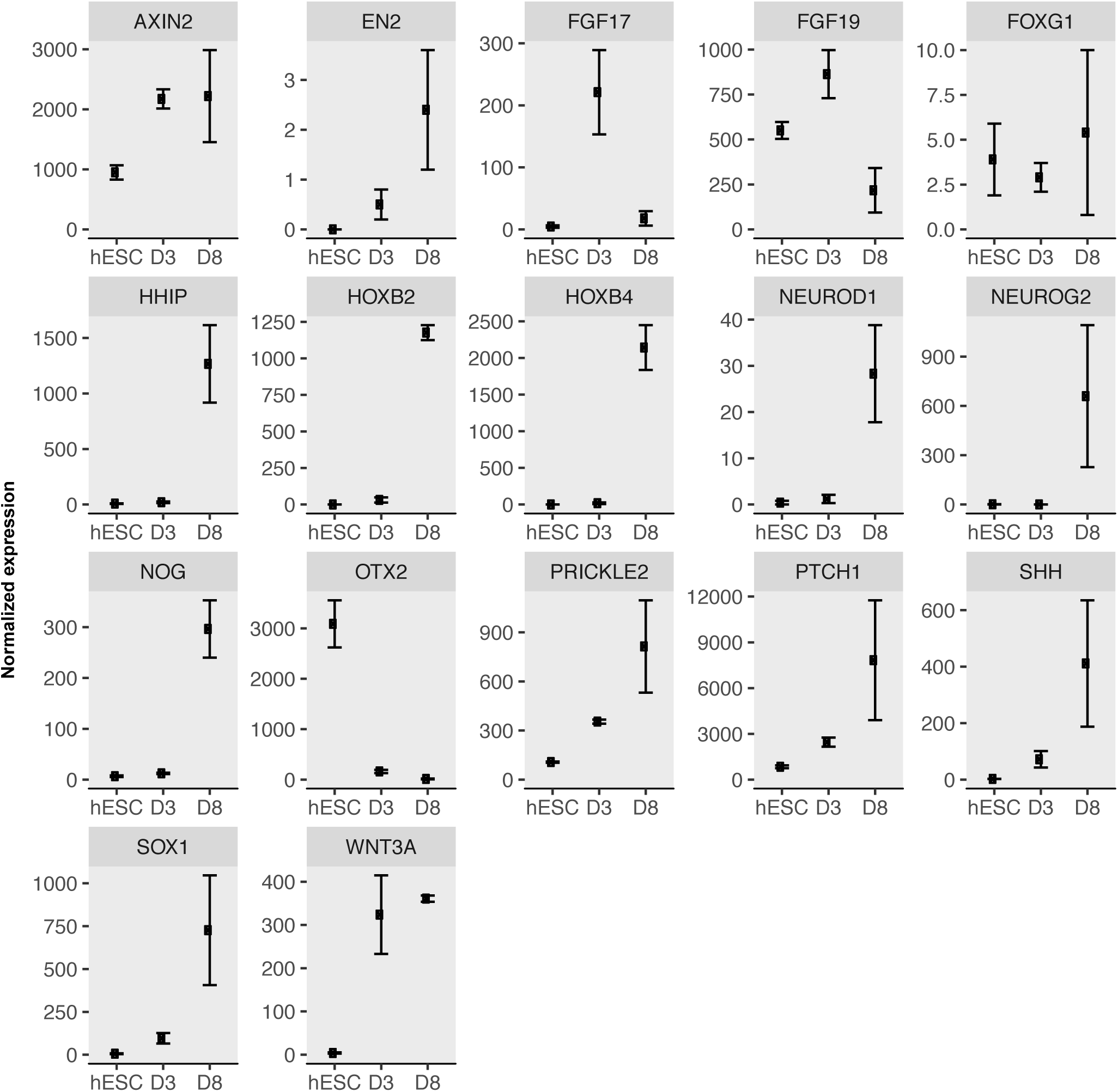
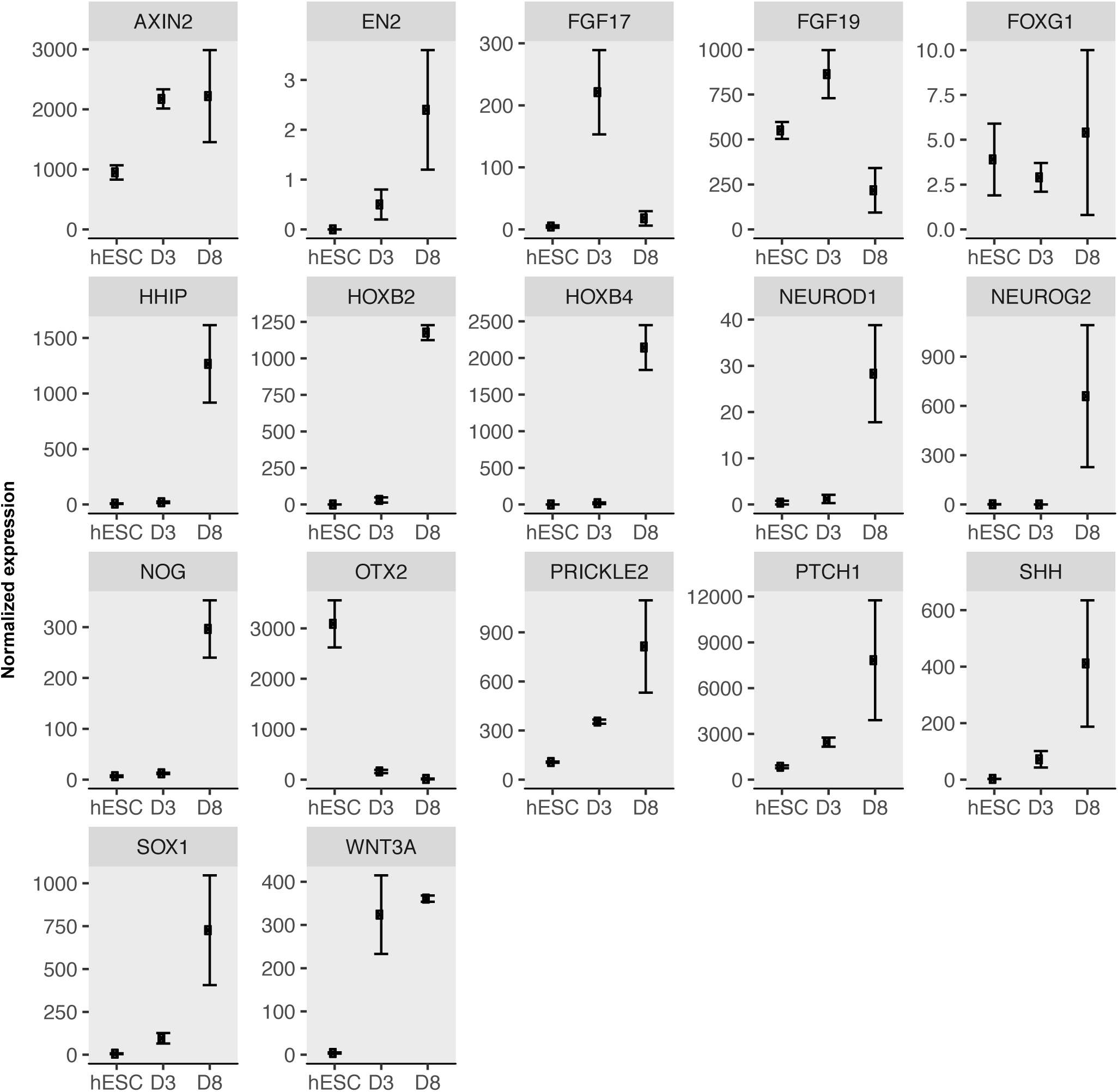
Standard Error Means for RNA-Seq data for each gene shown in Fig 4.

## References

Albano RM, Arkell R, Beddington RS, Smith JC. 1994. Expression of inhibin subunits and follistatin during postimplantation mouse development: decidual expression of activin and expression of follistatin in primitive streak, somites and hindbrain. Development 120: 803–813.

Andang M, Hjerling-Leffler J, Moliner A, Lundgren TK, Castelo-Branco G, Nanou E, Pozas E, Bryja V, Halliez S, Nishimaru H et al. 2008. Histone H2AX-dependent GABA(A) receptor regulation of stem cell proliferation. Nature 451: 460–464.

Ang S-L, Jin O, Rhinn M, Daigle N, Stevenson L, Rossant J. 1996. A targeted mouse Otx-2 mutation leads to severe defects in gastrulation and formation of axial mesoderm and to deletion of rostral brain. Development 122: 243–252.

Aulehla A, Wehrle C, Brand-Saberi B, Kemler R, Gossler A, Kanzler B, Herrmann BG. 2003. Wnt3a plays a major role in the segmentation clock controlling somitogenesis. Developmental cell 4: 395–406.

Beers J, Gulbranson DR, George N, Siniscalchi LI, Jones J, Thomson JA, Chen G. 2012. Passaging and colony expansion of human pluripotent stem cells by enzyme-free dissociation in chemically defined culture conditions. Nat Protoc 7: 2029–2040.

Bel-Vialar S, Medevielle F, Pituello F. 2007. The on/off of Pax6 controls the tempo of neuronal differentiation in the developing spinal cord. Developmental biology 305: 659–673.

Beverdam A, Meijlink F. 2001. Expression patterns of group-I aristaless-related genes during craniofacial and limb development. Mechanisms of development 107: 163–167.

Bulusu V, Prior N, Snaebjornsson MT, Kuehne A, Sonnen KF, Kress J, Stein F, Schultz C, Sauer U, Aulehla A. 2017. Spatiotemporal Analysis of a Glycolytic Activity Gradient Linked to Mouse Embryo Mesoderm Development. Developmental cell 40: 331–341 e334.

Chambers SM, Fasano CA, Papapetrou EP, Tomishima M, Sadelain M, Studer L. 2009. Highly efficient neural conversion of human ES and iPS cells by dual inhibition of SMAD signaling. Nature biotechnology 27: 275–280.

Chapman SC, Schubert FR, Schoenwolf GC, Lumsden A. 2002. Analysis of spatial and temporal gene expression patterns in blastula and gastrula stage chick embryos. Developmental biology 245: 187–199.

Cheng Y, Lotan R. 1998. Molecular cloning and characterization of a novel retinoic acid-inducible gene that encodes a putative G protein-coupled receptor. The Journal of biological chemistry 273: 35008–35015.

Chu LF, Leng N, Zhang J, Hou Z, Mamott D, Vereide DT, Choi J, Kendziorski C, Stewart R, Thomson JA. 2016. Single-cell RNA-seq reveals novel regulators of human embryonic stem cell differentiation to definitive endoderm. Genome Biol 17: 173.

Cunningham TJ, Brade T, Sandell LL, Lewandoski M, Trainor PA, Colas A, Mercola M, Duester G. 2015. Retinoic Acid Activity in Undifferentiated Neural Progenitors Is Sufficient to Fulfill Its Role in Restricting Fgf8 Expression for Somitogenesis. PloS one 10: e0137894.

Delfino-Machin M, Lunn JS, Breitkreuz DN, Akai J, Storey KG. 2005. Specification and maintenance of the spinal cord stem zone. Development 132: 4273–4283.

Denham M, Hasegawa K, Menheniott T, Rollo B, Zhang D, Hough S, Alshawaf A, Febbraro F, Ighaniyan S, Leung J et al. 2015. Multipotent caudal neural progenitors derived from human pluripotent stem cells that give rise to lineages of the central and peripheral nervous system. Stem cells 33: 1759–1770.

Diez del Corral R, Olivera-Martinez I, Goriely A, Gale E, Maden M, Storey K. 2003. Opposing FGF and retinoid pathways control ventral neural pattern, neuronal differentiation, and segmentation during body axis extension. Neuron 40: 65–79.

Gouti M, Delile J, Stamataki D, Wymeersch FJ, Huang Y, Kleinjung J, Wilson V, Briscoe J. 2017. A Gene Regulatory Network Balances Neural and Mesoderm Specification during Vertebrate Trunk Development. Developmental cell 41: 243–261 e247.

Gouti M, Metzis V, Briscoe J. 2015. The route to spinal cord cell types: a tale of signals and switches. Trends in genetics : TIG 31: 282–289.

Gouti M, Tsakiridis A, Wymeersch FJ, Huang Y, Kleinjung J, Wilson V, Briscoe J. 2014. In vitro generation of neuromesodermal progenitors reveals distinct roles for wnt signalling in the specification of spinal cord and paraxial mesoderm identity. PLoS biology 12: e1001937.

Harland R. 2000. Neural induction. Curr Opin Genet Dev 10: 357–362.

Hemmati Brivanlou A, Melton D. 1997. Vertebrate embryonic cells will become nerve cells unless told otherwise. Cell 88: 13–17.

Henrique D, Abranches E, Verrier L, Storey KG. 2015. Neuromesodermal progenitors and the making of the spinal cord. Development 142: 2864–2875.

Kim JH, Lee SR, Li LH, Park HJ, Park JH, Lee KY, Kim MK, Shin BA, Choi SY. 2011. High cleavage efficiency of a 2A peptide derived from porcine teschovirus-1 in human cell lines, zebrafish and mice. PloS one 6: e18556.

Komor AC, Badran AH, Liu DR. 2017. CRISPR-Based Technologies for the Manipulation of Eukaryotic Genomes. Cell 169: 559.

Kuroda H, Wessely O, De Robertis EM. 2004. Neural induction in Xenopus: requirement for ectodermal and endomesodermal signals via Chordin, Noggin, beta-Catenin, and Cerberus. PLoS biology 2: E92.

Liem KF, Jessell TM, Briscoe J. 2000. Regulation of the neural patterning activity of sonic hedgehog by secreted BMP inhibitors expressed by notochord and somites. Development 127: 4855–4866.

Linker C, Stern CD. 2004. Neural induction requires BMP inhibition only as a late step, and involves signals other than FGF and Wnt antagonists. Development 131: 5671–5681.

Lippmann ES, Williams CE, Ruhl DA, Estevez-Silva MC, Chapman ER, Coon JJ, Ashton RS. 2015. Deterministic HOX Patterning in Human Pluripotent Stem Cell-Derived Neuroectoderm. Stem cell reports 4: 632–644.

Liu H, Liu W, Maltby KM, Lan Y, Jiang R. 2006. Identification and developmental expression analysis of a novel homeobox gene closely linked to the mouse Twirler mutation. Gene expression patterns : GEP 6: 632–636.

Maddox DM, Condie BG. 2001. Dynamic expression of a glutamate decarboxylase gene in multiple non-neural tissues during mouse development. BMC Dev Biol 1: 1.

Meinhardt A, Eberle D, Tazaki A, Ranga A, Niesche M, Wilsch-Brauninger M, Stec A, Schackert G, Lutolf M, Tanaka EM. 2014. 3D reconstitution of the patterned neural tube from embryonic stem cells. Stem cell reports 3: 987–999.

Oginuma M, Moncuquet P, Xiong F, Karoly E, Chal J, Guevorkian K, Pourquie O. 2017. A Gradient of Glycolytic Activity Coordinates FGF and Wnt Signaling during Elongation of the Body Axis in Amniote Embryos. Developmental cell 40: 342–353 e310.

Olivera-Martinez I, Harada H, Halley PA, Storey KG. 2012. Loss of FGF-dependent mesoderm identity and rise of endogenous retinoid signalling determine cessation of body axis elongation. PLoS biology 10: e1001415.

Olivera-Martinez I, Storey KG. 2007. Wnt signals provide a timing mechanism for the FGF-retinoid differentiation switch during vertebrate body axis extension. Development 134: 2125–2135.

Padgett CL, Slesinger PA. 2010. GABAB receptor coupling to G-proteins and ion channels. Adv Pharmacol 58: 123–147.

Papanayotou C, De Almeida I, Liao P, Oliveira NM, Lu SQ, Kougioumtzidou E, Zhu L, Shaw A, Sheng G, Streit A et al. 2013. Calfacilitin is a calcium channel modulator essential for initiation of neural plate development. Nature communications 4: 1837.

Rausch T, Zichner T, Schlattl A, Stutz AM, Benes V, Korbel JO. 2012. DELLY: structural variant discovery by integrated paired-end and split-read analysis. Bioinformatics 28: i333–i339.

Ribes V, Stutzmann F, Bianchetti L, Guillemot F, Dolle P, Le Roux I. 2008. Combinatorial signalling controls Neurogenin2 expression at the onset of spinal neurogenesis. Developmental biology 321: 470–481.

Rodrigo-Albors A, Halley PA, Storey KG. 2016. Fate mapping caudal lateral epiblast reveals continuous contribution to neural and mesodermal lineages and the origin of the secondary neural tube. bioRxiv 045872.

Sasai N, Kutejova E, Briscoe J. 2014. Integration of Signals along Orthogonal Axes of the Vertebrate Neural Tube Controls Progenitor Competence and Increases Cell Diversity. PLoS biology 12: e1001907.

Scardigli R, Baumer N, Gruss P, Guillemot F, Le Roux I. 2003. Direct and concentration-dependent regulation of the proneural gene Neurogenin2 by Pax6. Development 130: 3269–3281.

Scardigli R, Schuurmans C, Gradwohl G, Guillemot F. 2001. Crossregulation between Neurogenin2 and pathways specifying neuronal identity in the spinal cord. Neuron 31: 203–217.

Schubert FR, Fainsod A, Gruenbaum Y, Gruss P. 1995. Expression of a novel murine homeobox gene Sax-1 in the developing nervous system. Mechanisms of development 51: 99–114.

Shull GE, Okunade G, Liu LH, Kozel P, Periasamy M, Lorenz JN, Prasad V. 2003. Physiological functions of plasma membrane and intracellular Ca2+ pumps revealed by analysis of null mutants. Ann N Y Acad Sci 986: 453–460.

Shum AS, Poon LL, Tang WW, Koide T, Chan BW, Leung YC, Shiroishi T, Copp AJ. 1999. Retinoic acid induces down-regulation of Wnt-3a, apoptosis and diversion of tail bud cells to a neural fate in the mouse embryo. Mechanisms of development 84: 17–30.

Sirbu IO, Duester G. 2006. Retinoic-acid signalling in node ectoderm and posterior neural plate directs left-right patterning of somitic mesoderm. Nature cell biology 8: 271–277.

Soldatov NM, Zuhlke RD, Bouron A, Reuter H. 1997. Molecular structures involved in L-type calcium channel inactivation. Role of the carboxyl-terminal region encoded by exons 40-42 in alpha1C subunit in the kinetics and Ca2+ dependence of inactivation. The Journal of biological chemistry 272: 3560–3566.

Spann P, Ginsburg M, Rangini Z, Fainsod A, Eyal Giladi H, Gruenbaum Y. 1994. The spatial and temporal dynamics of Sax1 (CHox3) homeobox gene expression in the chick's spinal cord. Development 120: 1817–1828.

Takemoto T, Uchikawa M, Kamachi Y, Kondoh H. 2006. Convergence of Wnt and FGF signals in the genesis of posterior neural plate through activation of the Sox2 enhancer N-1. Development 133: 297–306.

Tsakiridis A, Huang Y, Blin G, Skylaki S, Wymeersch F, Osorno R, Economou C, Karagianni E, Zhao S, Lowell S et al. 2014. Distinct Wnt-driven primitive streak-like populations reflect in vivo lineage precursors. Development 141: 1209–1221.

Tsakiridis A, Wilson V. 2015. Assessing the bipotency of in vitro-derived neuromesodermal progenitors. F1000Research 4: 100.

Turner DA, Hayward PC, Baillie-Johnson P, Rue P, Broome R, Faunes F, Martinez Arias A. 2014. Wnt/beta-catenin and FGF signalling direct the specification and maintenance of a neuromesodermal axial progenitor in ensembles of mouse embryonic stem cells. Development 141: 4243–4253.

Tzouanacou E, Wegener A, Wymeersch FJ, Wilson V, Nicolas JF. 2009. Redefining the progression of lineage segregations during mammalian embryogenesis by clonal analysis. Developmental cell 17: 365–376.

Wilson PA, Hemmati-Brivanlou A. 1995. Induction of epidermis and inhibition of neural fate by Bmp-4. Nature 376: 331–333.

Wilson V, Olivera-Martinez I, Storey KG. 2009. Stem cells, signals and vertebrate body axis extension. Development 136: 1591–1604.

Yamamoto A, Nagano T, Takehara S, Hibi M, Aizawa S. 2005. Shisa promotes head formation through the inhibition of receptor protein maturation for the caudalizing factors, Wnt and FGF. Cell 120: 223–235.

Young T, Deschamps J. 2009. Hox, Cdx, and anteroposterior patterning in the mouse embryo. Curr Top Dev Biol 88: 235–255.

Young T, Rowland JE, van de Ven C, Bialecka M, Novoa A, Carapuco M, van Nes J, de Graaff W, Duluc I, Freund JN et al. 2009. Cdx and Hox genes differentially regulate posterior axial growth in mammalian embryos. Developmental cell 17: 516–526.

